# Structure, receptor recognition and antigenicity of the human coronavirus CCoV-HuPn-2018 spike glycoprotein

**DOI:** 10.1101/2021.10.25.465646

**Authors:** M. Alejandra Tortorici, Alexandra C. Walls, Anshu Joshi, Young-Jun Park, Rachel T. Eguia, Terry Stevens-Ayers, Michael J. Boeckh, Amalio Telenti, Antonio Lanzavecchia, Davide Corti, Jesse D. Bloom, David Veesler

**Affiliations:** Department of Biochemistry, University of Washington, Seattle, WA 98195, USA; Basic Sciences Division, Fred Hutchinson Cancer Research Center, Seattle, WA 98109, USA; Vaccine and Infectious Disease Division, Fred Hutchinson Cancer Research Center, Seattle, WA 98109, USA; Vir Biotechnology, San Francisco, CA 94158, USA; Istituto Nazionale Genetica Molecolare, 20122 Milano, Italy; Humabs Biomed SA, a subsidiary of Vir Biotechnology, 6500 Bellinzona, Switzerland

## Abstract

The recent isolation of CCoV-HuPn-2018 from a child respiratory swab indicates that more coronaviruses are spilling over to humans than previously appreciated. Here, we determined cryo-electron microscopy structures of the CCoV-HuPn-2018 spike glycoprotein trimer in two distinct conformational states and identified that it binds canine, feline and porcine aminopeptidase N (APN encoded by *ANPEP*) orthologs which serve as entry receptors. Introduction of an oligosaccharide at position N739 of human APN renders cells susceptible to CCoV-HuPn-2018 spike-mediated entry, suggesting that single nucleotide polymorphisms could account for the detection of this virus in some individuals. Human polyclonal plasma antibodies elicited by HCoV-229E infection and a porcine coronavirus monoclonal antibody inhibit CCoV-HuPn-2018 S-mediated entry, indicating elicitation of cross-neutralizing activity among α-coronaviruses. These data provide a blueprint of the CCoV-HuPn-2018 infection machinery, unveil the viral entry receptor and pave the way for vaccine and therapeutic development targeting this zoonotic pathogen.

## Introduction

Four coronaviruses are endemic in humans and typically responsible for common colds: the β-coronaviruses OC43 and HKU1 and the α-coronaviruses NL63 and 229E. Moreover, three highly pathogenic β-coronaviruses have jumped from their animal hosts to humans in the last two decades: SARS-CoV, MERS-CoV and SARS-CoV-2. Both SARS-CoV and SARS-CoV-2 cluster in the sarbecovirus subgenus and originated in bats which most likely acted as reservoir hosts for these two viruses (Ge et al., 2013; Zhou et al., 2020). Whereas palm civets and racoon dogs have been recognized as intermediate hosts for zoonotic transmission of SARS-CoV between bats and humans (Guan et al., 2003), the SARS-CoV-2 intermediate host remains unknown. MERS-CoV was suggested to originate from bats although dromedary camels act as the reservoir host fueling spillover to humans (Haagmans et al., 2014; Memish et al., 2013). Recurrent coronavirus zoonoses along with detection of numerous coronaviruses in wildlife suggest that cross-species transmission events will continue to occur (Anthony et al., 2017; Menachery et al., 2015, 2016, 2020).

The coronavirus spike (S) glycoprotein folds as a homotrimer promoting viral entry into host cells (Tortorici and Veesler, 2019; Walls et al., 2016a, 2020a). S comprises an S_1_ subunit, which recognizes host cell receptors, and an S_2_ subunit, that mediates fusion of the viral and cellular membranes (Walls et al., 2017). Therefore, S plays a key role in modulating host and tissue tropism as well as zoonotic transmission. Since the coronavirus S glycoprotein is the target of neutralizing antibodies, it is the main focus of therapeutics and vaccine development, including monoclonal antibody therapies and vaccines (Baum et al., 2020; Corbett et al., 2020; Corti et al., 2021; Hansen et al., 2020; McCallum et al., 2021a; Piccoli et al., 2020; Pinto et al., 2020; Polack et al., 2020; Walls et al., 2019, 2020b, 2021).

Viruses genetically related to canine and feline coronaviruses have been previously identified in respiratory swabs obtained from patients with pneumonia or acute respiratory symptoms in Malaysia and the US (Silva and Mullis, 2014; Xiu et al., 2020). In Malaysia, eight) of 301 (2.5%) patients hospitalized with pneumonia between 2017 and 2018 in Sarawak were positive by pan-species coronavirus semi-nested RT-PCR assay and one specimen was identified as a novel canine-feline recombinant alphacoronavirus (genotype II) that was named CCoV-HuPn-2018 (Vlasova et al., 2021). The coronavirus isolate was shown to be cytopathic using canine A72 cells (Vlasova et al., 2021). These studies suggested that zoonotic transmission occurred and that these viruses, which were not previously known to infect humans, might be associated with clinical symptoms. Furthermore, these findings point to the circulation of several more coronaviruses in humans than previously appreciated and underscore the zoonotic threats posed by members of multiple distinct coronavirus genera.

### Architecture of the CCoV-HuPn-2018 S trimer

To visualize the infection machinery of the newly emerged CCoV-HuPn-2018, we characterized its S glycoprotein ectodomain trimer using cryo-electron microscopy (cryoEM). The CCoV-HuPn-2018 S trimer is ~150Å high with a triangular cross-section and comprises an N-terminal S_1_ subunit divided into domains designated 0 and A-D as well as a C-terminal S_2_ subunit which contains the fusion machinery (Tortorici and Veesler, 2019) **(Figure 1A)**. 3D classification of the data revealed the presence of two distinct S conformations with domain 0 swung out (at the periphery of the trimer) or proximal, i.e. oriented toward the viral membrane (apposed directly underneath domain A) for which we determined structures at 2.8 and 3.1 Å resolution, respectively **(Figure 1B-E, Figure S1 and Table S1)**. The latter domain 0 conformational state (‘proximal’) is similar to the one described for the S of HCoV-NL63 (Walls et al., 2016b) and porcine epidemic diarrhea virus (PEDV) (Kirchdoerfer et al., 2021) whereas the former conformation (‘swung out’) is more reminiscent of another state detected for PEDV S (Wrapp and McLellan, 2019) and feline infectious peritonitis virus (FIPV) **(Figure S2)**. We used local refinement to overcome the conformational variability of domain 0 and obtained reconstructions at 3.1 and 3.8 Å resolution for the swung out and proximal conformational states, respectively, which are related by a rotation of 135° **(Figure 1E, Figure S1 and Table S1)**.

**Figure 1.**
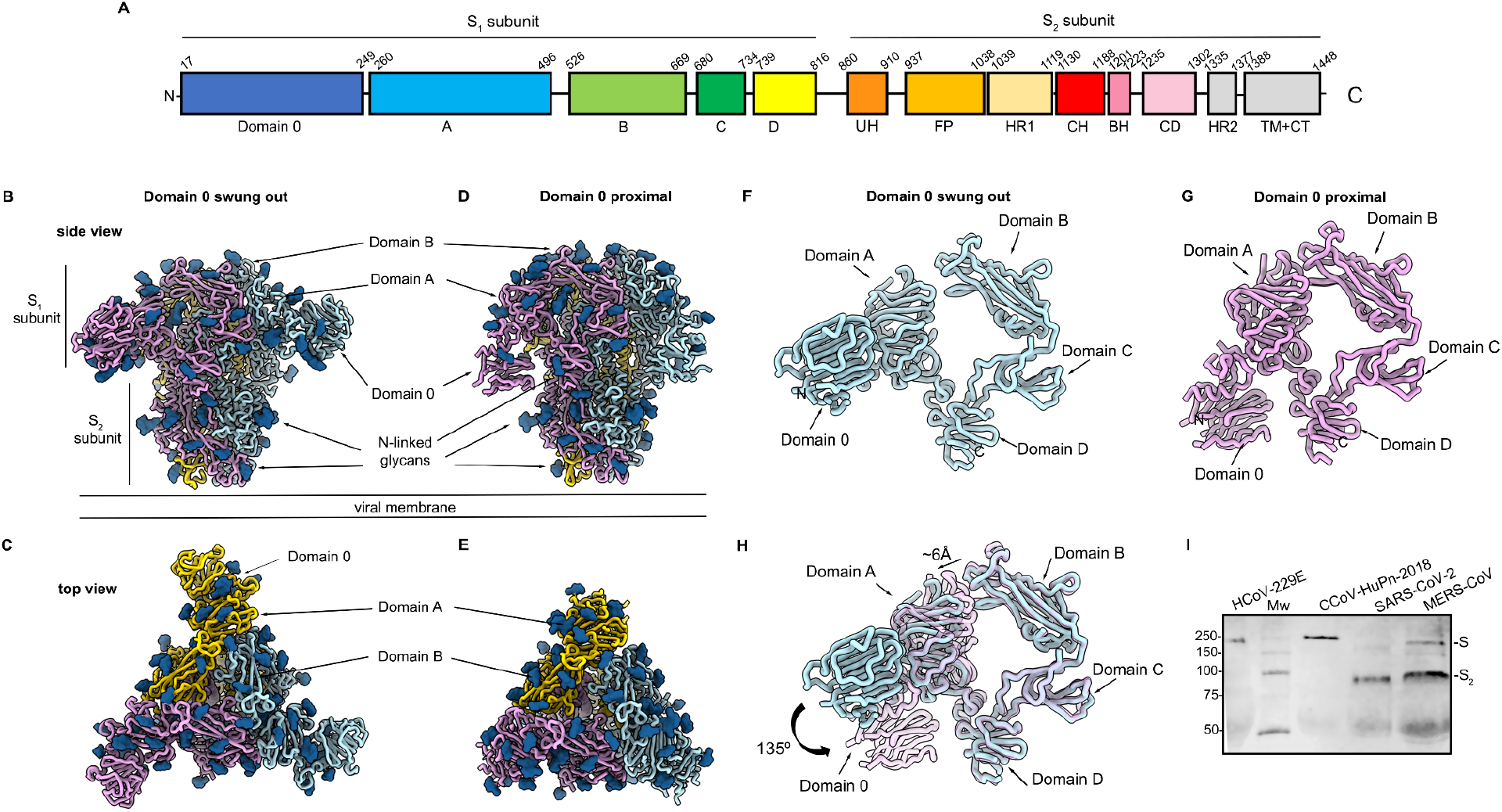
Architecture of the CCoV-HuPn-2018 infection machinery. **A,** Schematic diagram of the S glycoprotein organization. Dashed lines and grey boxes denote regions unresolved in the reconstruction and regions that were not part of the construct, respectively. UH, upstream helix; FP, fusion peptide; HR1, heptad-repeat 1; CH, central helix; BH, β-hairpin; CD: connector domain; HR2, heptad-repeat 2; TM, transmembrane domain; CT, cytoplasmic tail. **B-C,** CryoEM structure of the CCoV-HuPn-2018 S ectodomain trimer (with each domain 0 swung out) viewed along two orthogonal orientations. **D-E,** CryoEM structure of the CCoV-HuPn-2018 S ectodomain trimer (with each domain 0 pointing ‘proximal’ toward the viral membrane) viewed along two orthogonal orientations. **F-G,** Ribbon diagram of the CCoV-HuPn-2018 S_1_ subunit with domain 0 in the swung out (F) and proximal (G) conformations. **H,** Superimposition of the S_1_ subunit from the structures with domain 0 in the swung out (light blue) and proximal (pink) conformations showing the A domain moving away from the B domain of the same protomer and rotation of domain 0. **I,** Western blot of VSV pseudotyped particles harboring HCoV-229E S, CCoV-HuPn-2018 S, SARS-CoV-2 S or MERS-CoV S. Mw: molecular weight ladder. Full-length S and S_2_ subunit bands are indicated on the right hand side of the blot.

The CCoV-HuPn-2018 S trimer is densely decorated with oligosaccharides distributed in both the S_1_ and S_2_ subunits. In total, 24 glycans out of 32 putative N-linked glycosylation sequons are resolved to different extents in at least one of the two cryoEM maps **(Figure 1B-E and Table S2)**. The S_1_ subunit structure is most similar to NL63 and 229E S whereas the S_2_ subunit architecture is most closely related to PEDV and FIPV S, supporting the possibility that the CCoV-HuPn-2018 S gene arose from a recombination event (Vlasova et al., 2021). Although the B domains remain in a closed state, i.e. sitting atop the fusion machinery, in both CCoV-HuPn-2018 S conformations, the two structures differ in the tertiary and quaternary organization of the S_1_ subunit crown. Specifically, domain A moves radially outwards upon transition from a proximal to a swung out domain 0 conformation, positioning it >6Å further away from the B domain belonging to the same protomer **(Figure 1F-H)**. As a result, domain A moves closer to and forms tight interactions with the C domain of a neighboring protomer in the swung out domain 0 state. Overall, interactions among C domains contribute to a marked enhancement of the total surface area buried between S_1_ protomers (stabilizing the S_1_ trimeric trimer) whereas the B domains reduce tertiary and quaternary contacts with their surroundings, likely enabling subsequent opening and receptor engagement (cf. below).

The CCoV-HuPn-2018 S lacks a polybasic furin cleavage site at the S_1_/S_2_ junction, similar to type II feline and canine coronaviruses (Licitra et al., 2013; Millet and Whittaker, 2015). Accordingly, we did not detect proteolytic processing of CCoV-HuPn-2018 S transfected in HEK293T cells, similar to HCoV-229E S **(Figure 1G)**. In contrast, transfection of HEK293T cells with SARS-CoV-2 S or MERS-CoV S leads to cleavage at S_1_/S_2_ by furin-like proteases **(Figure 1G)** (Millet and Whittaker, 2014, 2015; Walls et al., 2020a). The CCoV-HuPn-2018 S2’ site contains a polybasic motif K953RKYR957 which is identical to the one found in TGEV S and some type II feline and canine coronavirus S sequences (Millet and Whittaker, 2015) and might modulate cleavage and infectivity.

### Organization of the CCoV-HuPn-2018 S_1_ subunit

Coronaviruses can use multiple domains within the S_1_ subunit for receptor attachment and some coronavirus spike glycoproteins engage more than one receptor type to enter host cells (Hulswit et al., 2019; Lempp et al., 2021; Li et al., 2017; Park et al., 2019; Peng et al., 2011; Tortorici et al., 2019). Domain A folds as a galectin-like β-sandwich and most closely resembles the equivalent domains of α-coronavirus S glycoproteins including HCoV-NL63 (Walls et al., 2016b), HCoV-229E (Li et al., 2019), PEDV (Kirchdoerfer et al., 2021; Wrapp and McLellan, 2019) and FIPV (Hsu et al., 2020) with which it shares ~40% sequence identity. Superimposition of the CoV-HuPn-2018 S 0 and A domains underscores their conserved architecture and topology (r.m.s.d. 4.2 Å over 139 aligned Cα positions) although they share only 12.5% sequence identity **(Figure 2 A-G)**. These findings suggest that both domains derived from an ancestral gene duplication event, as previously described for HCoV-NL63 (Walls et al., 2016b), and accumulated extensive mutations leading to their very low sequence relatedness.

**Figure 2.**
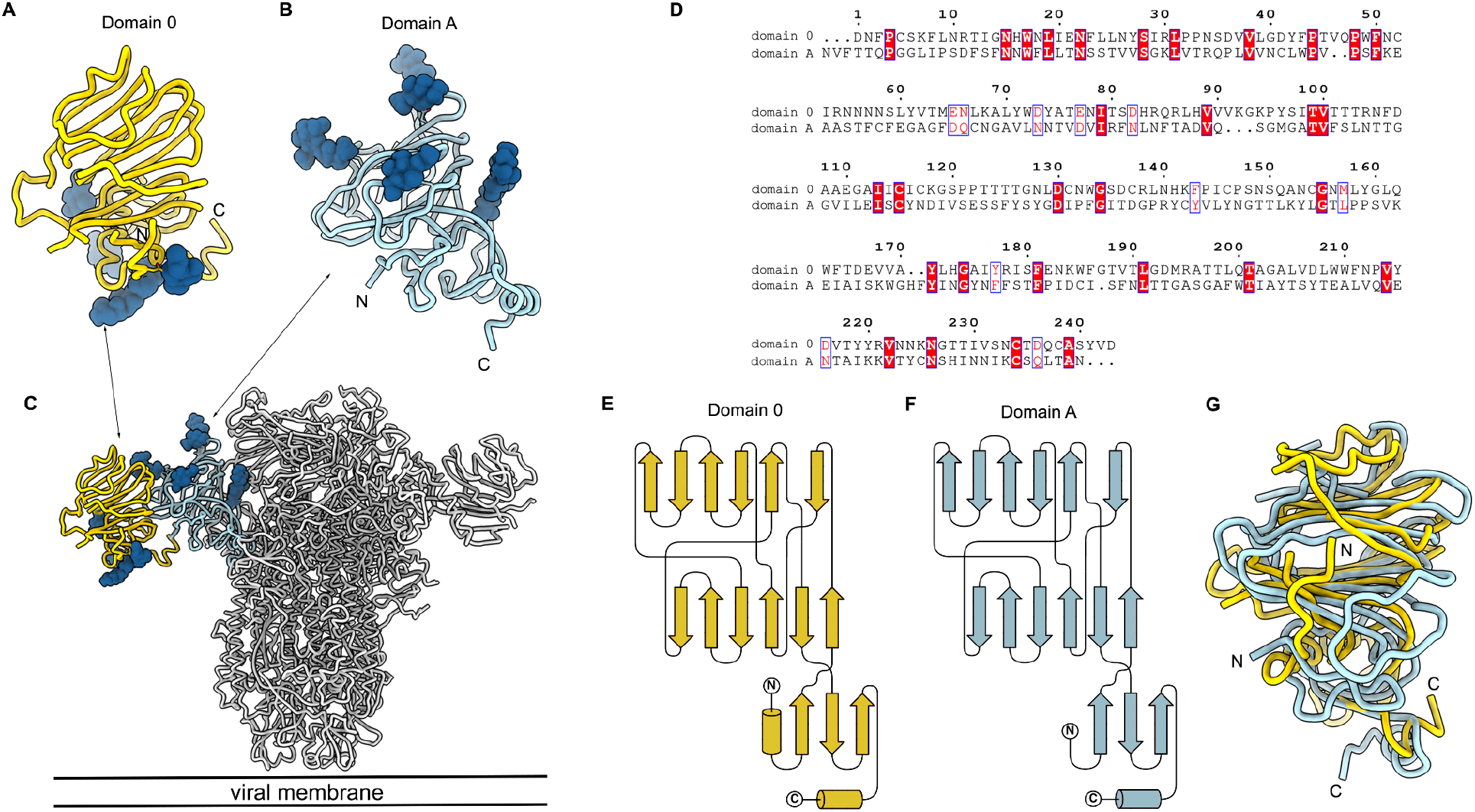
Structural conservation of CCoV-HuPn-2018 S domain 0 and domain A. **A-C,** Ribbon diagrams of the CCoV-HuPn-2018 S domain 0 (A) and domain A (B) oriented as shown in the prefusion S conformation shown in (C). **D,** Sequence alignment of CCoV-HuPn-2018 S domain A and domain 0 highlighting the low sequence identity (12.5%) between them. **E-G,** Topology diagrams of domain 0 (E) and domain A (F) and structural overlay between the two domains (G) underscoring their similarity. N-linked glycans are rendered as blue spheres in panels (A-C) but were removed from panel G for clarity.

The CCoV-HuPn-2018 domain B structure is strikingly similar to the porcine respiratory coronavirus (PRCV) and transmissible gastroenteritis virus (TGEV) domain B structures (Reguera et al., 2012) with which it shares ~90% sequence identity, including key APN interacting residues (PRCV Y528/W571 corresponding to CCoV-HuPn-2018 Y543/W586) **(Figure 3A-C)**. Although the β-sandwich fold is also shared with HCoV-NL63, HCoV-229E and porcine delta-coronavirus, their receptor-binding loops adopt distinct conformations compared to CCoV-HuPn-2018, PRCV and TGEV (Reguera et al., 2012; Shang et al., 2018; Wong et al., 2017; Wu et al., 2009; Xiong et al., 2018) **(Figure S3).**

**Figure 3.**
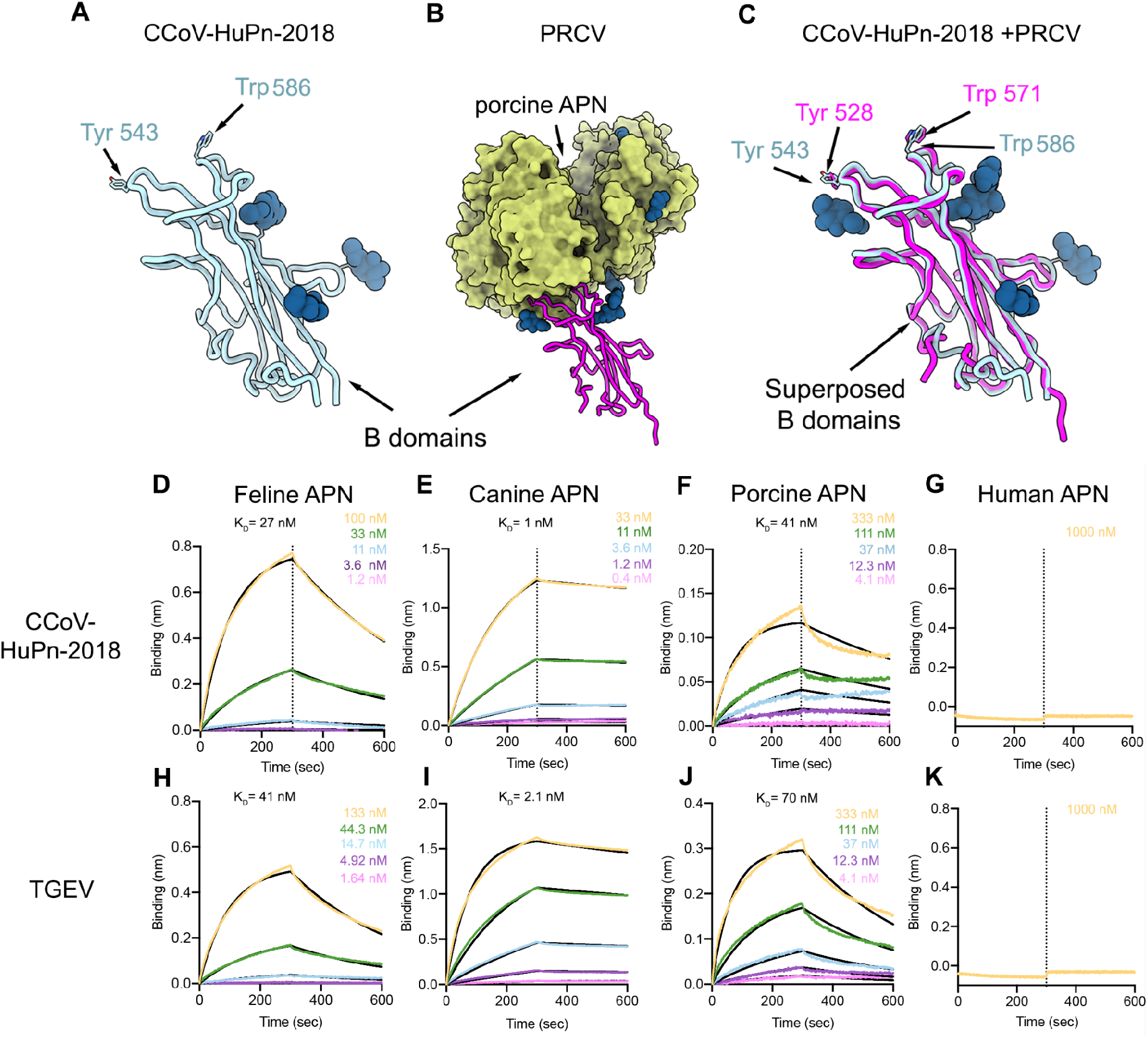
The CCoV-HuPn-2018 S B domain recognizes APN. **A-C,** Ribbon diagrams of the CCoV-HuPn-2018 B domain (A), the PRCV B domain bound to porcine APN (PDB: 4F5C) (B) and overlay (C) underscoring their structural similarity. N-linked glycans are rendered as blue spheres. **D-K,** Biolayer interferometry kinetic binding analysis of feline (D,H), canine (E,I), porcine (F,J) and human (G,K) monomeric APN ectodomains to biotinylated CCoV-HuPn-2018 B domain (D-G) or TGEV B domain (H-K) immobilized at the surface of SA biosensors.

### Aminopeptidase N is a receptor for CCoV-HuPn-2018

Based on the structural similarity between the B domain of CCoV-HuPn-2018 **(Figure 3A-C)**, and other alpha CoVs such as PRCV and TGEV, we evaluated the ability of the CCoV-HuPn-2018 B domain to interact with several APN orthologs which is a receptor for both PRCV and TGEV (Delmas et al., 1992). Feline, canine and porcine APNs, but not human APN, recognized biotinylated CCoV-HuPn-2018 B domain immobilized at the surface of biolayer interferometry biosensors **(Figure 3D-G)**. The CCoV-HuPn-2018 B domain bound with higher affinity to canine APN (K_D_=1.0 nM) compared to feline APN (K_D_=27nM) and porcine APN (K_D_=41nM), as was the case for the TGEV B domain **(Figure 3D-K, Table S3)**. We also evaluated the ability of the CCoV-HuPn-2018 B domain to interact with several APN orthologs by pull-down assays and found that feline, canine and porcine APNs, but not human APN, recognized biotinylated CCoV-HuPn-2018 and TGEV B domain coupled to streptavidin beads **(Figure S4A-B,D-E)**. Human APN interacted with the HCoV-229E B domain as observed by pull-down and BLI **(Figure S4C-G and Table 3)**. These data suggest that CCoV-HuPn-2018 can use several APN orthologs as receptors likely involving a binding mode comparable to PRCV/TGEV but distinct from HCoV-229E (Reguera et al., 2012; Wong et al., 2017).

We next assessed the ability of various APN orthologs to promote cell entry of vesicular stomatitis virus (VSV) particles pseudotyped with CCoV-HuPn-2018 S, TGEV S or HCoV-229E S. Transient transfection of feline, canine or porcine - but not human-APN, rendered HEK293T cells susceptible to CCoV-HuPn-2018 S VSV pseudovirus entry **(Figure 4A)**, in line with the binding data (**Figure 3D-G**). Feline, canine and porcine APNs also promoted TGEV S pseudotyped virus entry into HEK293T cells **(Figure 4B)** whereas HCoV-229E S-mediated entry occurred with feline or human APN but not with canine or porcine orthologs **(Figure 4C)**. Cotransfection of TMPRSS2 with APN did not enhance CCoV-HuPn-2018 S pseudovirus entry **(Figure 4D-F)**, likely reflecting the ability to employ redundant proteolytic pathways for membrane fusion to occur at the plasma membrane or the endosomal membrane, similar to HCoV-229E (Bertram et al., 2013). We also observed CCoV-HuPn-2018 S VSV entry in canine A72 cells (Vlasova et al., 2021), canine MDCK cells, and feline CRFK cells (**Figure S5),** presumably due to endogenous APN expression. Among the cell lines evaluated, feline and canine cell lines permitted TGEV S VSV entry whereas only feline CRFK cells facilitated entry of 229E S pseudovirus (**Figure S5)**, in line with the data obtained with transfected APN orthologs (**Figure 4A-C).** Finally, we observed concentration-dependent inhibition of CCoV-HuPn-2018 S pseudotyped virus entry using purified dimeric feline, canine or porcine APN-Fc fusions **(Figure 4G)**, establishing APN as a bona fide entry receptor for CCoV-HuPn-2018.

**Figure 4.**
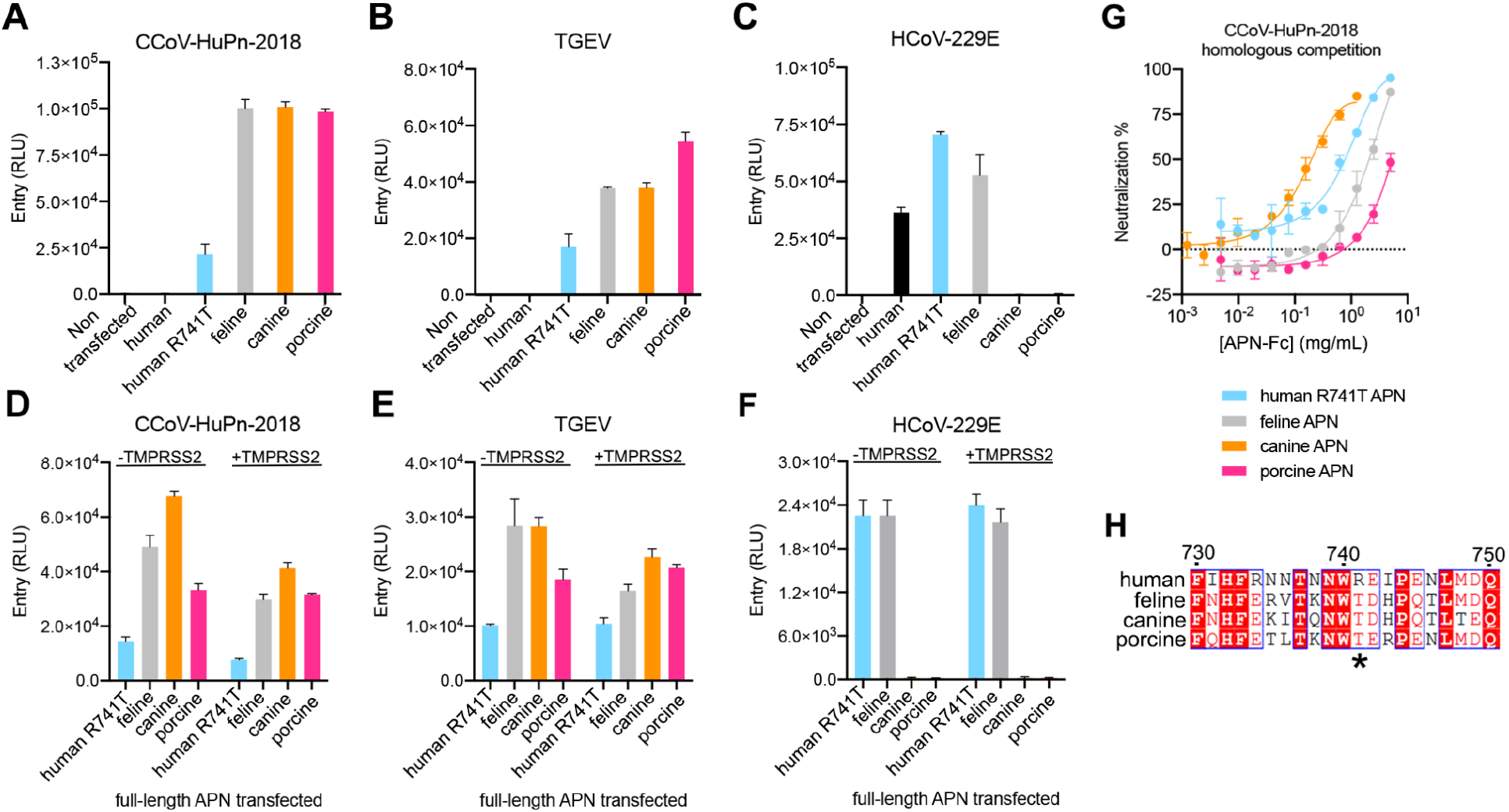
APN is a functional entry receptor for CCoV-HuPn-2018. **A-C,** Entry of VSV particles pseudotyped with the CCoV-HuPn-2018 S (A), TGEV S (B) and HCoV-V229E S (C) in HEK293T cells transiently transfected with canine, feline, porcine, human or human R741T (glycan knockin) APN orthologs. RLU: relative luciferase units. **D-F**, Entry of VSV particles pseudotyped with the CCoV-HuPn-2018 S (D), TGEV S (E) and HCoV-V229E S (F) in HEK293T cells transiently transfected with human R741T (glycan knockin), canine, feline, or porcine APN orthologs with or without TMPRSS2. **E,** Concentration-dependent inhibition of CCoV-HuPn-2018 S pseudovirus entry in HEK293T cells transiently transfected with full-length APN orthologs with matched, purified dimeric soluble APN-Fc ectodomains. **H,** Sequence alignment of human, feline, canine, and porcine APNs focused on the N739 glycosylation sequon. Human APN position 741 is indicated with an asterisk.

A key distinction between human APN and feline/canine/porcine APNs is the absence of an N-linked oligosaccharide at position N739 of the former orthologue due to a Thr to Arg residue substitution at position 741, which is position i+2 of a glycosylation sequon (**Figure 4H**). Previous work showed that abrogation of this glycosylation site prevented binding of the TGEV RBD to cellsurface expressed porcine APN (Reguera et al., 2012; Tusell et al., 2007). The structural conservation of the CCoV-HuPn-2018 and TGEV/PRCV B domains suggest that the absence of this oligosaccharide is responsible for the lack of detectable binding and inability of CCoV-HuPn-2018 S to utilize human APN for cell entry. To evaluate this hypothesis, we generated a human APN N739 oligosaccharide knockin mutant (by introducing the R741T substitution) and assessed its ability to mediate pseudotyped virus entry. Human R741T APN rendered HEK293T cells permissive to both CCoV-HuPn-2018 and TGEV S-mediated entry (**Figure 4A,B**) and human R741T APN-Fc inhibited CCoV-HuPn-2018 S pseudovirus entry in a concentration-dependent manner (**Figure 4G**), confirming the key role of this post-translational modification for receptor recognition. Moreover, the human APN oligosaccharide knockin mutant remained functional as demonstrated by its retained interactions with the HCoV-229E B domain **(Figure S4F-G)** and its ability to support 229E S-mediated entry into HEK293T cells (**Figure 4C**) due to the distal location of the N739 oligosaccharide from the HCoV-229E attachment site (Wong et al., 2017).

### Antigenicity of the CCoV-HuPn-2018 S trimer

CCoV-HuPn-2018 S shares 47% amino acid sequence identity with HCoV-229E S and with HCoV-NL63 S and is therefore more distantly related to the S glycoproteins of these two endemic human-infecting α-coronaviruses than they are to each other (64% sequence identity). Mapping sequence conservation between these three viruses on the CCoV-HuPn-2018 S trimer structure shows that the S_1_ subunit is more divergent than the S_2_ subunit (**Figure 5A-B**), a trend which holds true across all coronaviruses as a result of differential receptor usage and immune pressure (Tortorici and Veesler, 2019; Walls et al., 2020a).

**Figure 5.**
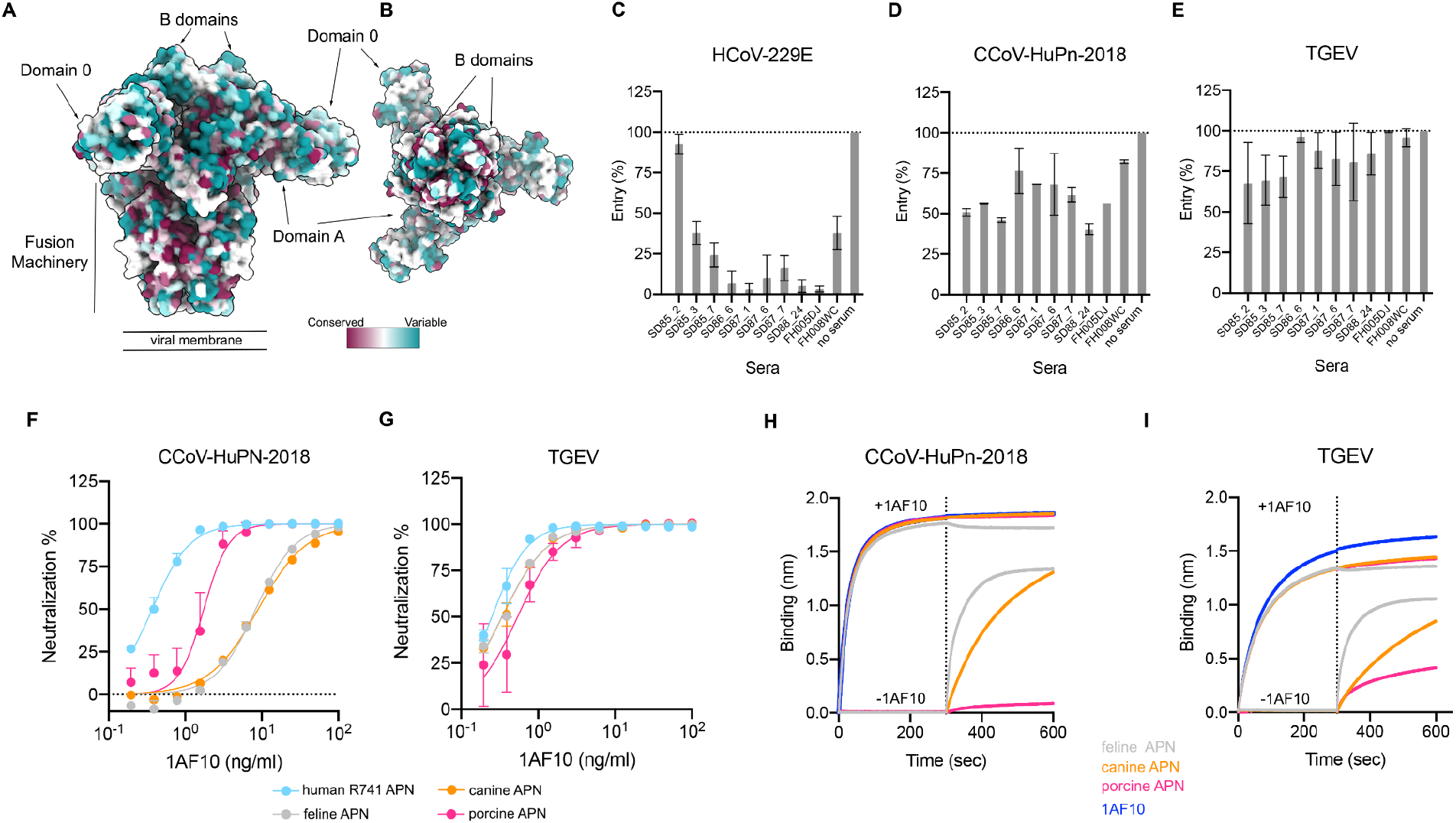
Evaluation of polyclonal and monoclonal antibody neutralization of CCoV-HuPn-2018 S-mediated entry. **A-B**, Sequence conservation of HCoV-229E, HCoV-NL63 and CCoV-HuPn-2018 S glycoproteins plotted on the CCoV-HuPn-2018 S structure viewed from the side (A) and top (B). The sequence alignment was generated using the following sequences: CCoV-HuPn-2018 S (QVL91811.1), HCoV-229E S (AAK32191.1 and ABB90515) and HCoV-NL63 S (AIW52835.1 and YP_003767.1). **C-E,** CCoV-HuPn-2018 S (C), TGEV S (D) and HCoV-229E S (E) pseudotyped virus entry in the presence of a 1:5 diluted plasma obtained between 1985 and 1987 from human subjects previously infected with HCoV-229E (Full neutralization curves are shown in Figure S6). **F-G** CCoV-HuPn-2018 S (F) and TGEV S (G) pseudotyped virus entry in the presence of various concentrations of 1AF10 neutralizing monoclonal Fab in HEK293T cells transfected with the indicated full-length APN orthologs. **H-I,** Competition between 1AF10 Fab and APN for binding to immobilized CoV-HuPn-2018 (H) or TGEV (I) B domains analyzed by biolayer interferometry. Each CoV-HuPn-2018 (H) or TGEV (I) B domain-loaded SA biosensor was sequentially dipped in a solution containing 10X above the measured affinity (Table S3) with 170 or 100 nM 1AF10 Fab respectively and then 170 or 100 nM 1AF10 Fab + 10 or 21 nM canine (orange), 270 or 470nM feline (grey), 410 or 700nM porcine (pink) APN monomeric or 170 or 100 nM 1AF10 Fab alone (blue).

To investigate the CCoV-HuPn-2018 cross-neutralization afforded by prior α-coronavirus exposure, we evaluated the serum neutralizing activity of human plasma obtained between 1985 and 1987 from individuals infected with HCoV-229E (Eguia et al., 2021). We observed that plasma inhibiting HCoV-229E S-mediated entry into cells also blocked CCoV-HuPn-2018 S pseudovirus entry, albeit with reduced potency (**Figure 5C-D and Figure S6**). In contrast, TGEV S-mediated entry was little affected by any plasma tested (**Figure 5E and Figure S6**). These findings suggest that α-coronavirus infection in humans elicit polyclonal antibody responses with some degree of neutralization breadth towards other coronaviruses in the same genera.

We subsequently evaluated the ability of the previously described PRCV/TGEV 1AF10 neutralizing monoclonal antibody (Wong et al., 2017) to cross-react with the CCoV-HuPn-2018 B domain. Biolayer interferometry measurements showed that the 1AF10 Fab fragment bound to the immobilized CCoV-HuPn-2018 B domain with an affinity of 17 nM with similar on and off rates than those observed for binding to the TGEV B domain **(Figure S7)**. Moreover, the 1AF10 Fab potently inhibited CCoV-HuPn-2018 S and TGEV S pseudovirus entry into HEK293T cells transiently transfected with canine, feline, porcine and human APN N739 oligosaccharide knockin mutant (R741T) **(Figure 5F-G)**. We also demonstrated that 1AF10 inhibits APN binding competitively to both the CCoV-HuPn-2018 and the TGEV B domains indicating that this mAb prevents viral attachment to the host cell surface and would be a suitable therapeutic candidate against CCoV-HuPn-2018 **(Figure 5H-I)**.

## Discussion

Since several other α-coronaviruses interacts with cell surface carbohydrates (Krempl et al., 1997; Milewska et al., 2014; Regan and Whittaker, 2008; Regan et al., 2010), we suggest that the two snapshots we captured using cryoEM correspond to two functionally distinct conformations at various stages of the entry process. These structural changes would couple domain 0 adsorption to host cell surface glycans (putative attachment receptors) to subsequent exposure of the B domain to interact with the APN entry receptor. Subsequent protease-mediated cleavage at the S2’ site activates the spike for membrane fusion and enables initiation of infection. As similar conformational changes were detected for PEDV S which also bind glycans using domain 0 (Kirchdoerfer et al., 2021; Wrapp and McLellan, 2019), we propose that this cascade of conformational changes might be conserved among several α-coronaviruses.

We previously described that NL63 S domain A N358 glycan participates in obstructing the B domain receptor-binding loops from the same protomer as a putative immune evasion strategy (Walls et al., 2016b) (**Figure S8A**). The CCoV-HuPn-2018 S domain A glycan at position N404 and the domain B glycan at position N561 respectively shield the receptor-binding loops from the same and from a neighboring B domain (to a different extent in the domain 0 proximal and swung out conformations) **(Figure S8C-D)**, and both might therefore play a similar role for this virus. Both CCoV-HuPn-2018 S and HCoV-NL63 S therefore appear to utilize conformational masking and glycan shielding to limit exposure of the receptor-binding loops and possibly reduce recognition by neutralizing antibodies (Reguera et al., 2012; Walls et al., 2016b; Wong et al., 2017).

The ongoing SARS-CoV-2 genetic drift has led to the emergence of variants harboring numerous mutations, especially in the N-terminal domain (domain A) and the receptor-binding domain (domain B) (Collier et al., 2021; Davies et al., 2021; Deng et al., 2021; Faria et al., 2021; McCallum et al., 2021a, 2021b; Tegally et al., 2021; Thomson et al., 2021). Many of these variants exhibit altered transmissibility, immune evasion, replication kinetics or disease severity relative to the ancestral SARS-CoV-2 isolate (Cele et al., 2021; Collier et al., 2021; Davies et al., 2021; Edara et al., 2021; Liu et al., 2021a, 2021b; McCallum et al., 2021a, 2021b; Plante et al., 2020; Wibmer et al., 2021). Similarly, a longitudinal survey of HCoV-229E isolates revealed that the (B domain) receptor-binding loops evolve rapidly relative to the rest of the genome resulting in changes in receptor-binding affinity and antigenicity (Wong et al., 2017). This antigenic drift results in dampened serum neutralization titers of HCoV-229E isolates that have emerged several years after infection (Eguia et al., 2021). Nevertheless, the existence in HCoV-229E patient sera of some cross-reactive Abs with detectable neutralizing activity against CCoV-HuPn-2018 has the potential to reduce disease severity of this virus in humans.

Single nucleotide polymorphisms of the APN gene (*ANPEP*) have been previously described in humans (Vijgen et al., 2004) and pigs (Bovo et al., 2021) and we identified a rare R741G-carrying allele in a heterozygous individual in public databases (cf. methods), which does not, however, introduce a glycosylation sequon. Moreover, we observed a near-perfect sequence conservation of the CCoV-HuPn-2018 receptor-binding loops with those of TGEV/PRCV although they are distinct viruses. Collectively, these data along with the failure of CCoV-HuPn-2018 S to utilize wildtype human APN and its isolation using canine A72 cells (Vlasova et al., 2021), suggest that CCoV-HuPn-2018 is not well adapted for infection and replication in human hosts. These findings also suggest that patients infected with this novel virus or with related canine and feline coronaviruses (Silva and Mullis, 2014; Vlasova et al., 2021) might harbor *ANPEP* variants introducing an oligosaccharide at position N739 which we have shown to be necessary and sufficient to support CCoV-HuPn-2018 S-mediated entry. The rarity of these single nucleotide polymorphisms is expected to restrict the zoonotic potential of CCoV-HuPn-2018 to a small number of individuals in the absence of further viral adaptations.

The detection of viruses genetically related to canine and feline coronaviruses, including CCoV-HuPn-2018, in humans across two different continents revealed that cross-species transmission of α-coronavirus-1 species, which were not previously known to infect humans, might occur recurrently (Silva and Mullis, 2014; Vlasova et al., 2021; Xiu et al., 2020). Moreover, demonstration of the ability of porcine deltacoronavirus to use human APN for cell entry and identification of multiple zoonotic transmission events in children in Haiti add an additional coronavirus to the growing list of emerging human pathogens (Lednicky et al., 2021; Li et al., 2018). The data presented here along with continued surveillance and characterization of emerging coronaviruses will support pandemic preparedness efforts similar to prior work that enabled a rapid response to the COVID-19 pandemic.

## Author contributions

M.A.T. and D.V. conceived the project. M.A.T., A.C.W and D.V. designed experiments. A.C.W. and A.J. performed BLI binding assays. M.A.T. and A.J. carried out pull-down assays. M.A.T. ran pseudotyped virus entry assays. M.A.T. and A.J. expressed and purified the recombinant proteins. M.A.T. and Y.J.P. collected the cryoEM data. M.A.T. processed the cryoEM data. M.A.T. and D.V. built and refined the atomic models. A.T. performed the SNP analysis. R.T.E., T.S.A., M.J.B. A.L., D.C. and J.B. provided unique reagents. M.A.T. and D.V. wrote the manuscript with input from all authors.

## Acknowledgements

We thank Hideki Tani (University of Toyama) for providing the reagents necessary for preparing VSV pseudotyped viruses. This study was supported by the National Institute of Allergy and Infectious Diseases (DP1AI158186 and HHSN272201700059C to D.V.), the National Institute of General Medical Sciences (R01GM120553 to D.V.), a Pew Biomedical Scholars Award (D.V.), an Investigators in the Pathogenesis of Infectious Disease Awards from the Burroughs Wellcome Fund (D.V.), Fast Grants (D.V.) and the Bill & Melinda Gates Foundation (OPP1156262 to D.V.), the University of Washington Arnold and Mabel Beckman cryoEM center and the National Institute of Health grant S10OD032290 (to D.V.).

---

**Figure S1.**
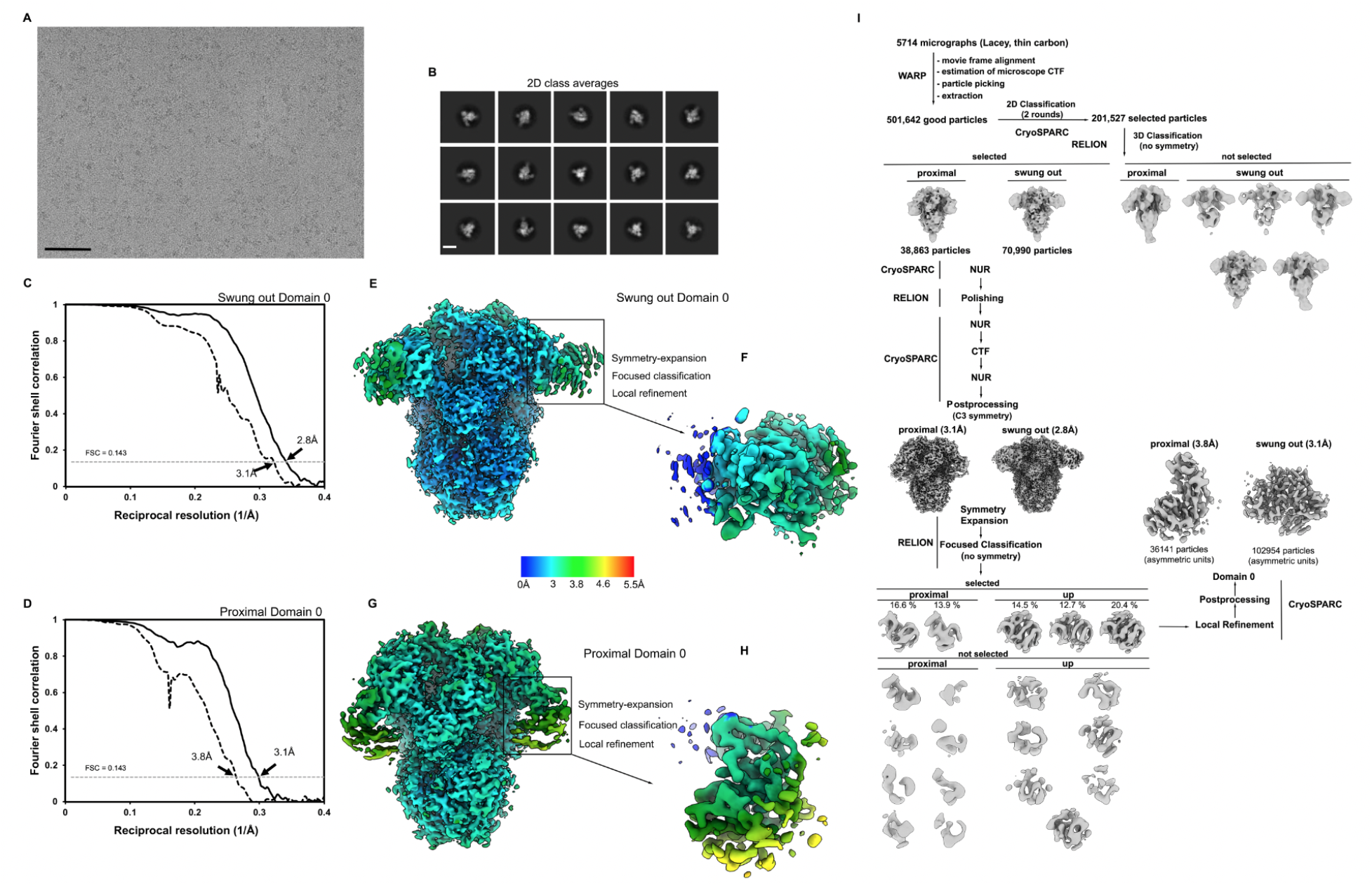
CryoEM data processing and validation CCoV-HuPn-2018 S dataset. **A-B,** Representative electron micrograph (A) and class averages (B) of CCoV-HuPn-2018 S in complex with the S2X259 Fab. Scale bar of the micrograph: 500 Å. Scale bar of the class averages: 100 Å. **C-D,** Gold-standard Fourier shell correlation curves for the CCoV-HuPn-2018 S trimer (solid black line) and the locally refined reconstruction of the CCoV-HuPn-2018 S domain 0 (dashed black line) from the swung domain 0 conformation (C) and the down domain 0 conformation (D). The 0.143 cut-off is indicated by a horizontal dashed grey line. **E-H,** Local resolution map for the CCoV-HuPn-2018 S trimer and the locally refined reconstruction of the domain 0, in the swung conformation (E,F) and in the down conformation (G, H). I, CryoEM data processing flow-chart.

**Figure S2.**
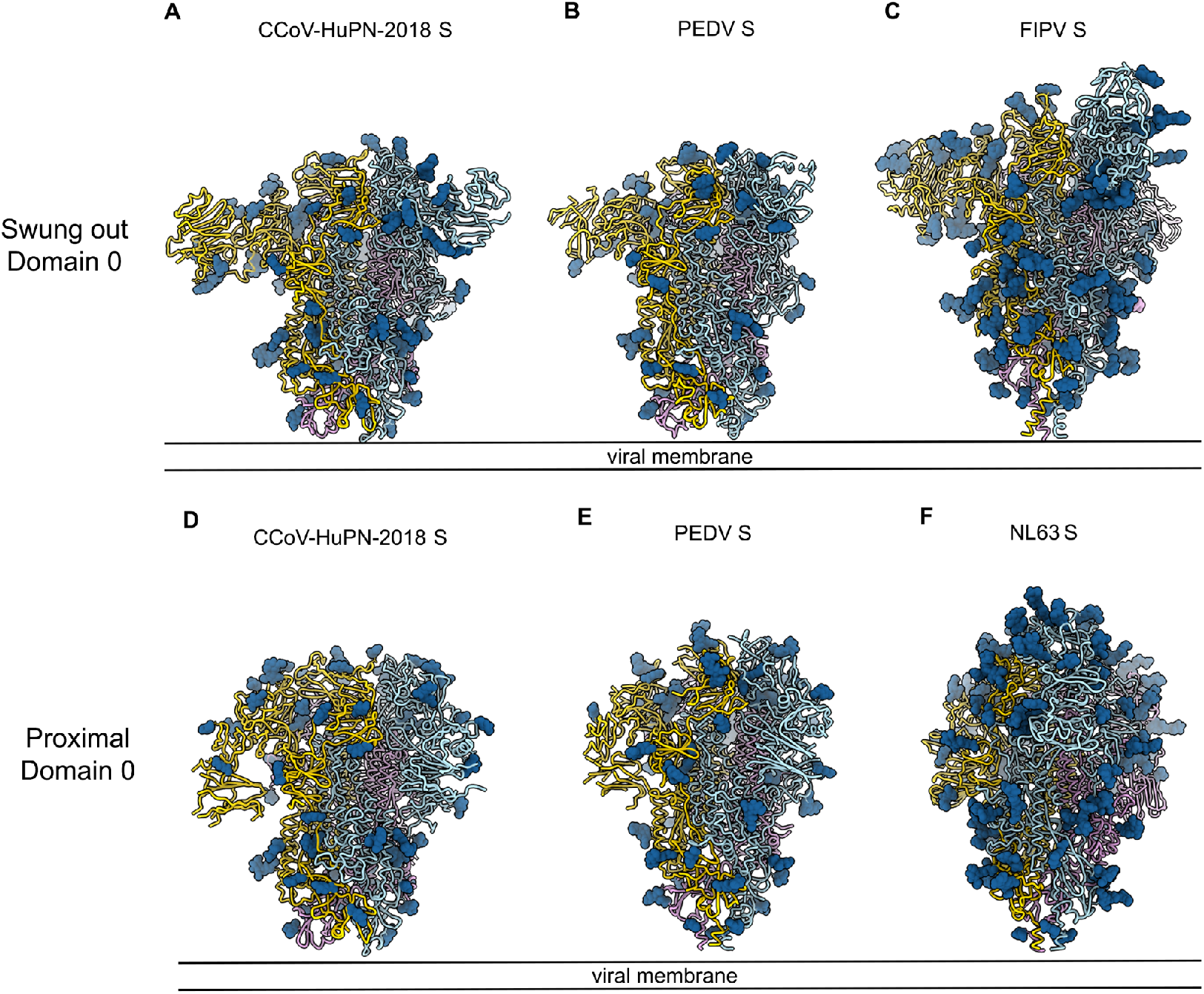
Architecture of α-coronavirus S trimers harboring a domain 0 obtained by Cryo-EM. **A-F,** Side view of S trimers from CCoV-HuPn-2018 with the two conformations of domain 0 (A, D). PEDV S (PDB 6VV5) (B), FIPV S (PDB 6JX7) (C), PEDV S (PDB 6U7K) (E), NL63 S (PDB 5SZS) (F). Each α-coronavirus S protomer is colored distinctly (light blue, pink and gold). S trimers are rendered in ribbon representation. N-linked glycans are shown as blue spheres.

**Figure S3.**
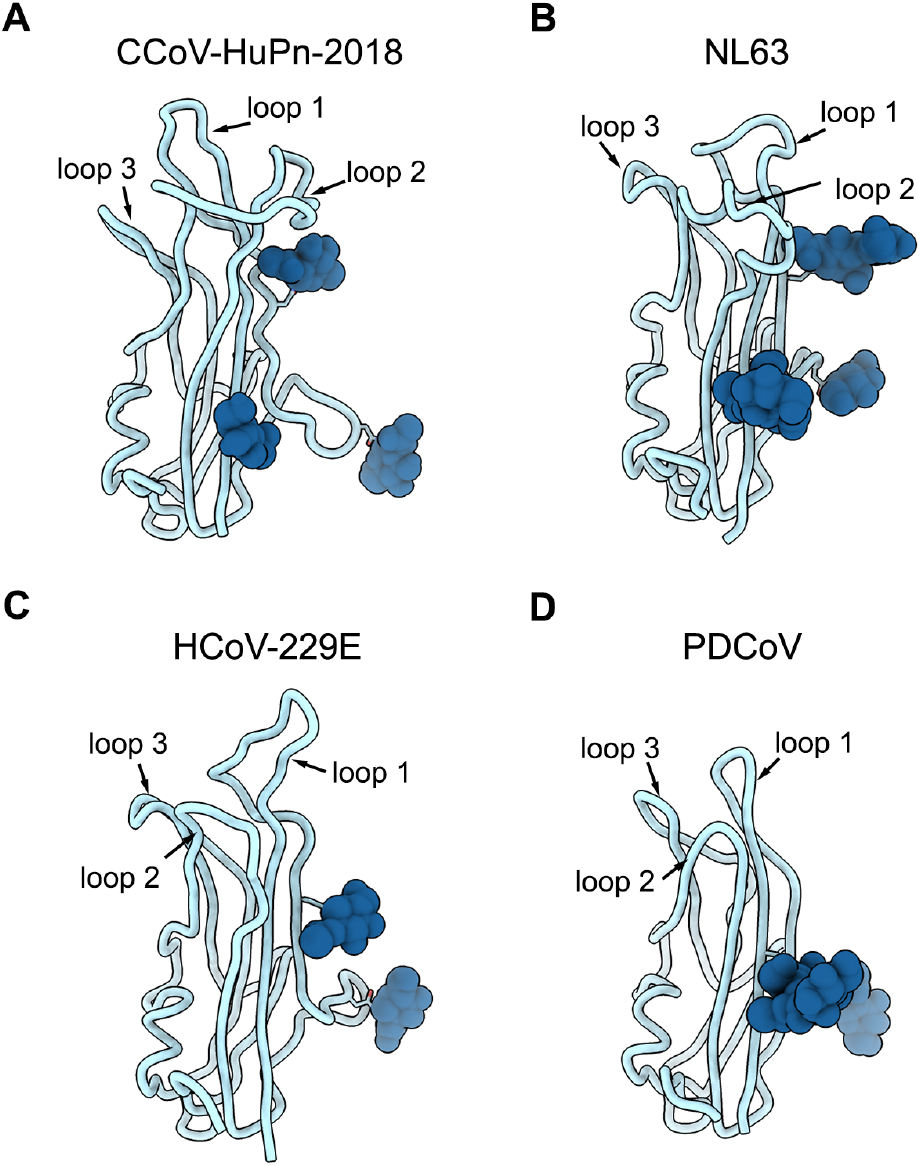
Architecture of α-coronavirus B domains. **A-D,** Ribbon diagrams of B domains from CCoV-HuPn-2018 (A), NL63 (PDB 5SZS) (Walls et al., 2016b) (B), HCoV-229E (PDB 6ATK) (Wong et al., 2017) (C) and PDCoV (PDB 6BFU) (Xiong et al., 2018) (D). B domains from NL63, HCoV229-E and PDCoV were structurally aligned with the B domain from CCoV-HuPn-2018. All B domains are colored light blue and N-linked glycans are rendered as blue spheres. The receptor binding loops are indicated.

**Figure S4.**
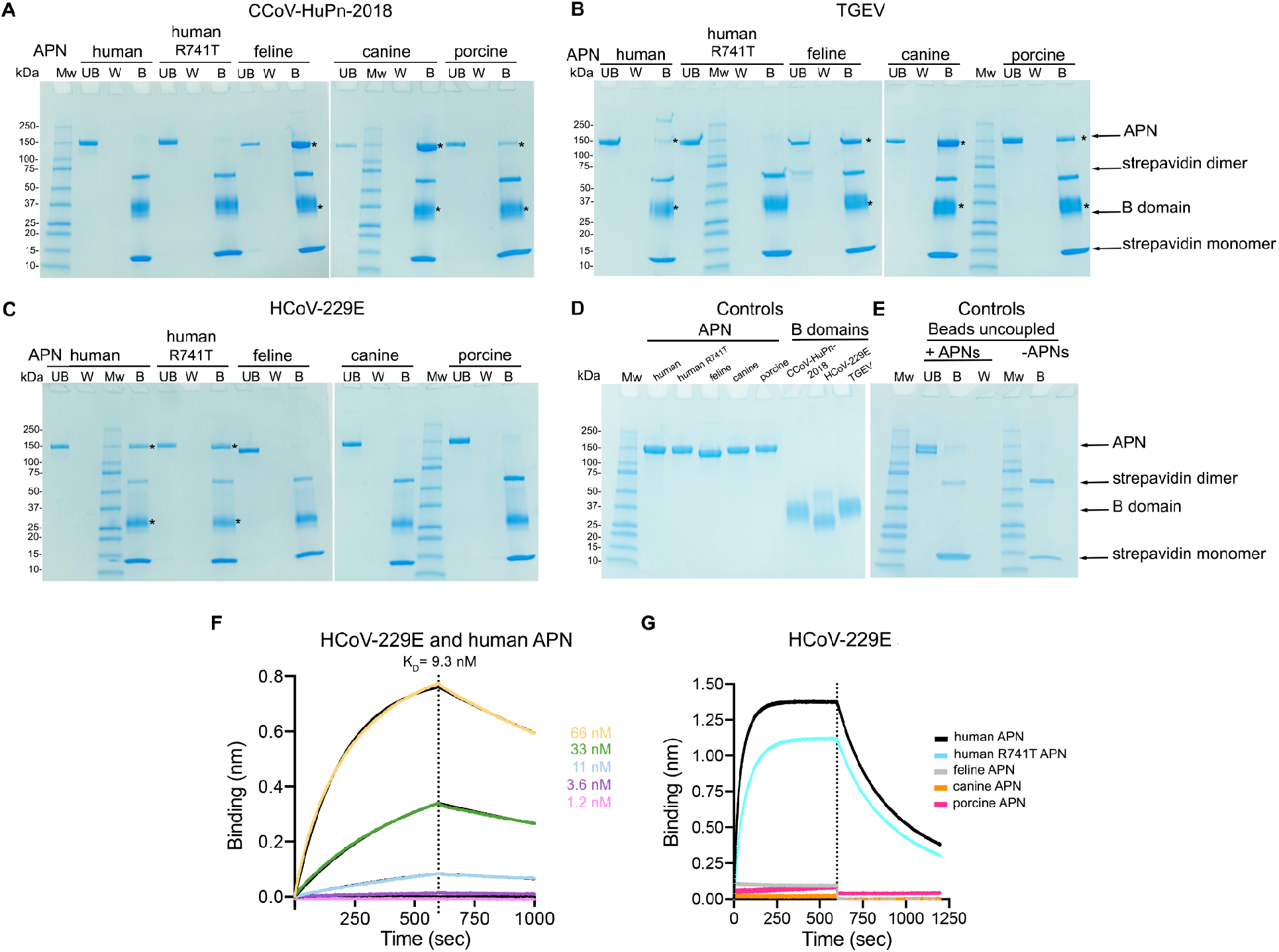
The CCoV-HuPn-2018 B domain recognizes APN orthologs. Coomassie stained SDS-PAGE analysis of the interaction between biotinylated B domains from CCoV-HuPn-2018 (A), TGEV (B) and HCoV-229E (C) immobilized on magnetic streptavidin beads and APN-Fc orthologs assessed by pull-down assays. Asterisks indicate the positions of the APN/B domain complex pulled-down. UB: unbound W: wash, B: bound. Purified proteins used in the pull-down assay (D) and beads without B domain (uncoupled) incubated or not with APNs analyzed by Coomassie stained SDS-PAGE (E). F. Biolayer interferometry kinetic binding analysis of human monomeric APN ectodomains to biotinylated HCoV-229E immobilized at the surface of SA biosensors. G. Biolayer interferometry kinetic binding analysis of human, human R741T, feline, canine and porcine monomeric APN ectodomains at 1 μM to biotinylated HCoV-229E immobilized at the surface of SA biosensors.

**Figure S5.**
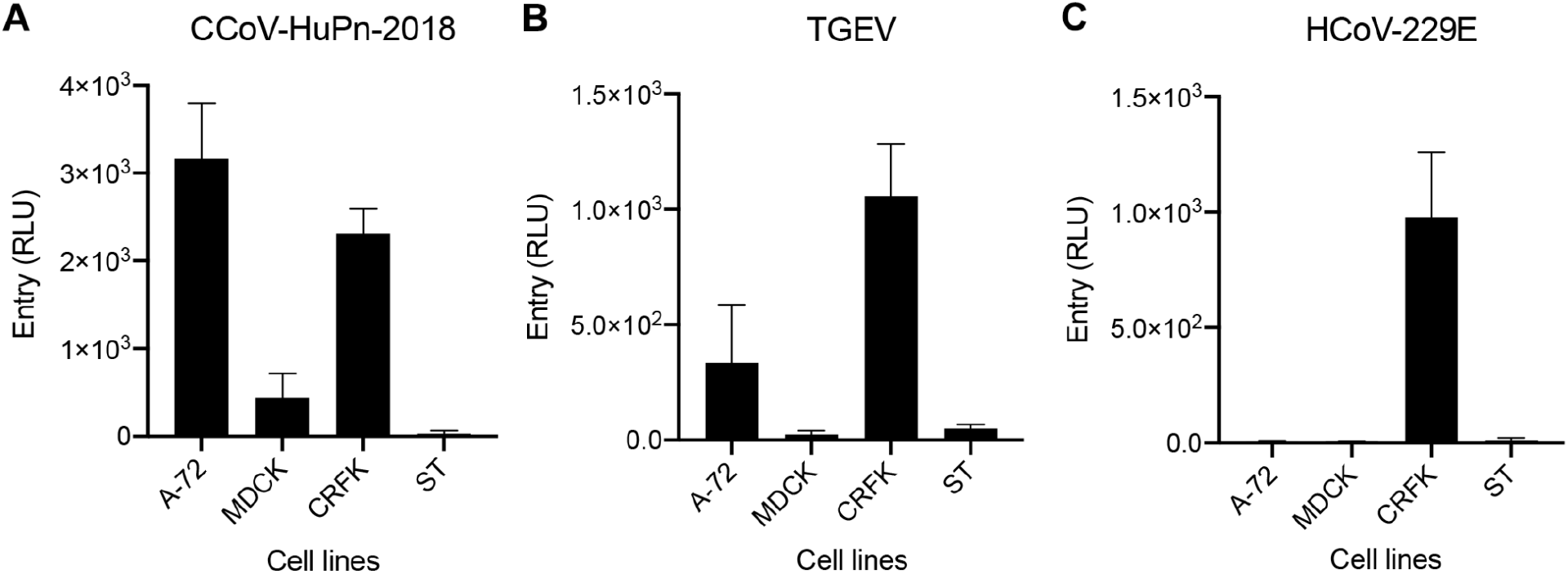
Evaluation of pseudotyped virus mediated entry in feline, canine and pig cell lines. A-C. CCoV-HuPn-2018 S (A), TGEV S (B) and HCoV-229E S (C) pseudotyped virus mediated entry in *Canis familiaris* tumor fibroblast cells (A-72), *Canis familiaris* epithelial kidney cells (MDCK), *Felis catus* (CRFK) and *Sus scrofa* pig testis fibroblast cells (ST).

**Figure S6.**
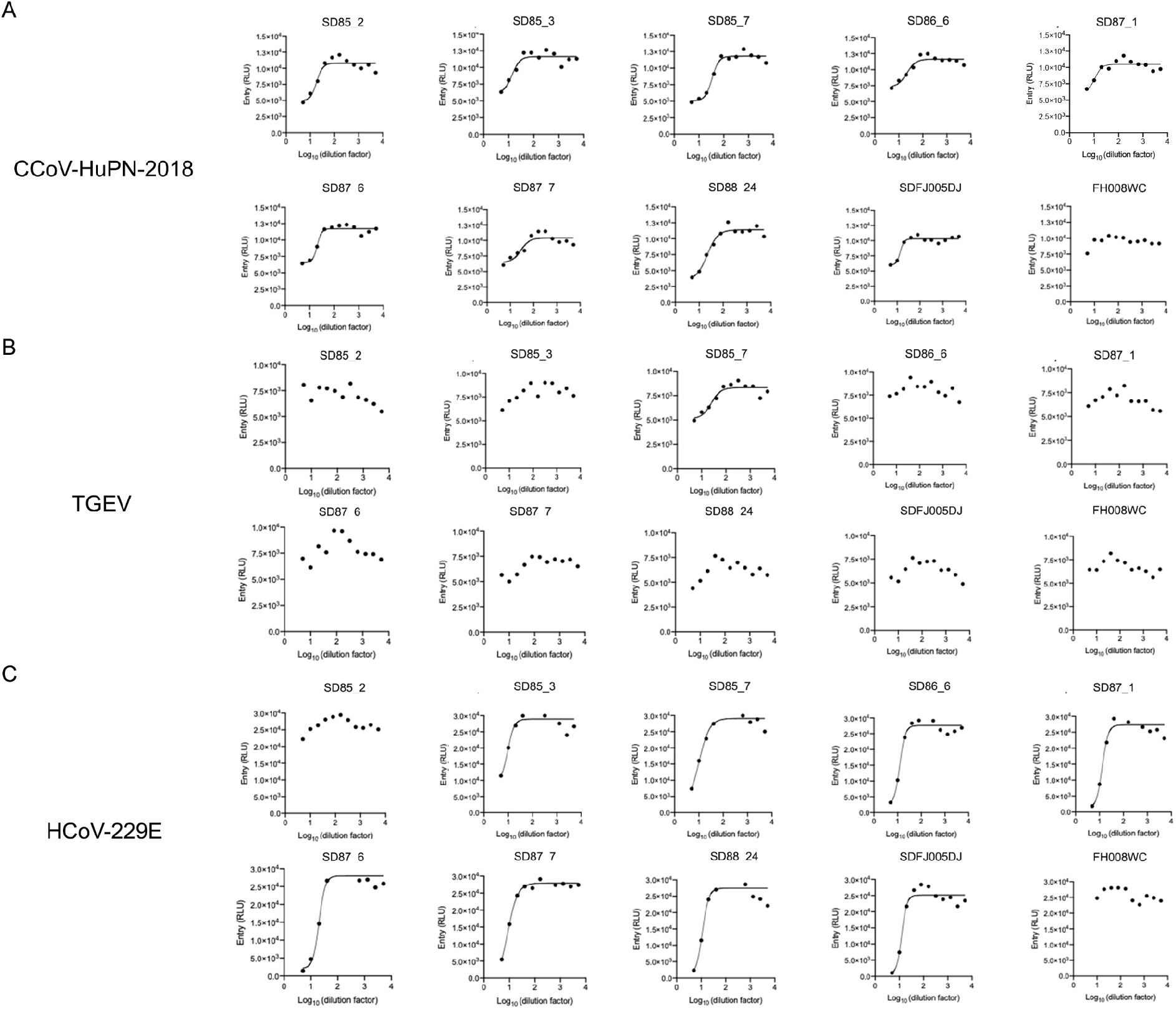
Evaluation of sera neutralization of CCoV-HuPn-2018 S-mediated entry. **A-C,** CCoV-HuPn-2018 (A), TGEV (B) and HCoV-229E (C) S pseudotyped virus mediated entry in the presence of various dilutions of plasma (name of the sample is indicated at the top of each graph) obtained between 1985 and 1989 from human subjects previously infected with HCoV-229E. Shown is one representative experiment out of two. Fits are shown only when inhibition of entry was observed.

**Figure S7.**
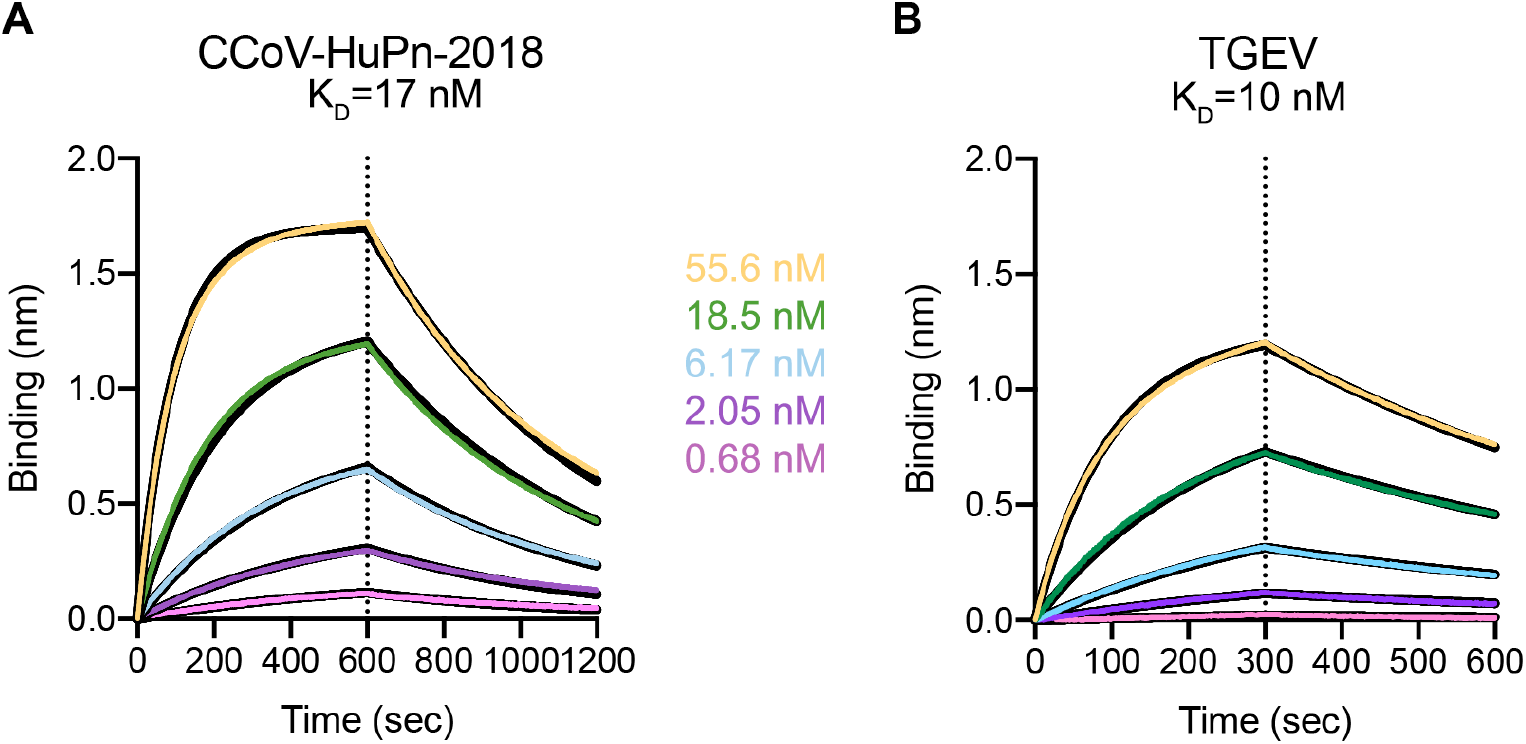
Binding affinity for monoclonal Fab 1AF10. Biolayer interferometry kinetic binding analysis of 1AF10 Fab to biotinylated CCoV-HuPn-2018 (A) and TGEV (B) B domains immobilized at the surface of SA biosensors.

**Figure S8.**
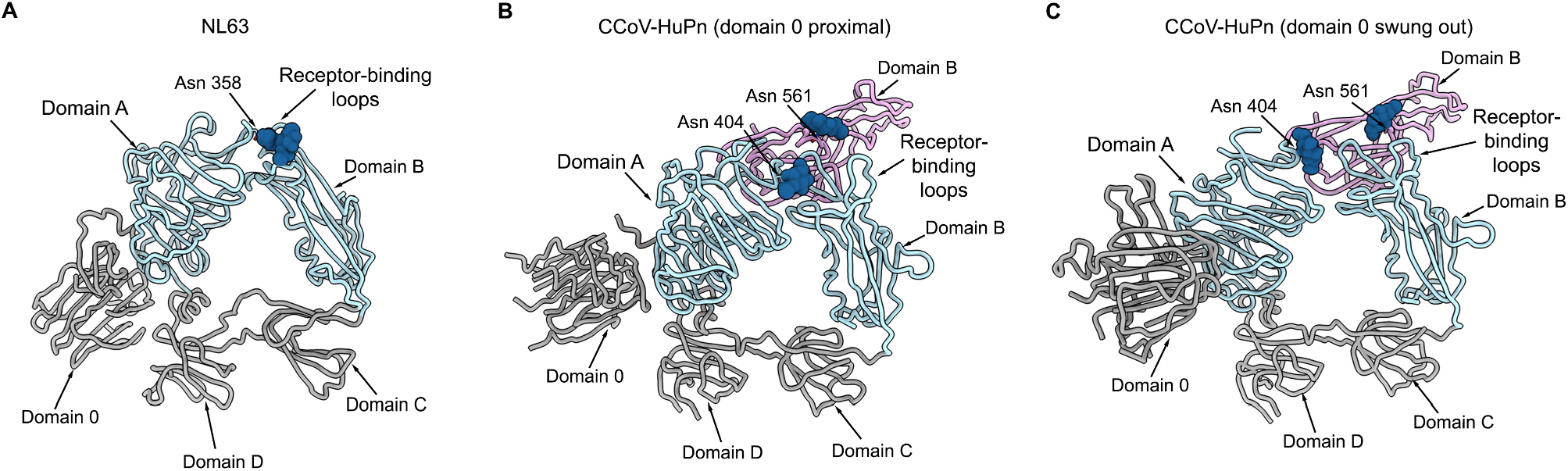
Glycan-mediated immune-evasion strategy of alphacoronavirus S trimers. Ribbon diagrams of S_1_ subunits of NL63 (PDB 5SZS) (A) and CCoV-HuPn-2018 with domain 0 in proximal conformation (B) and domain B in swung out conformation (C). The HCoV-NL63 receptor-binding loops are buried through interactions with domain A of the same protomer (light blue), including the glycan moiety at Asn358, and are not available to engage host-cell receptors (A). The CCoV-HuPn-2018 receptor-binding loops are also buried by interactions with domain A, including the glycan moiety at Asn404 of the same protomer (light blue), but also the glycan moiety at Asn561 from another protomer (pink) (B and C).

**Table S1.** CryoEM data collection and refinement statistics for CCoV-HuPn-2018 S with domain 0 in swung out and proximal conformations.

|  | S trimer<br>(swung out) | Domain 0<br>(swung out, local<br>refinement) | S trimer<br>(proximal) | Domain 0<br>(proximal, local<br>refinement) |
| --- | --- | --- | --- | --- |
| **Data collection and**<br>**processing** |  |  |  |  |
| Magnification | 130,000 | 130,000 | 130,000 | 130,000 |
| Voltage | 300 | 300 | 300 | 300 |
| Electron exposure (e-/Å<sup>2</sup>) | 84 | 84 | 84 | 84 |
| Defocus range (μm) | -0.8/-2.0 | -0.8/-2.0 | -0.8/-2.0 | -0.8/-2.0 |
| Pixel size (Å) | 0.8430 | 0.8430 | 0.8430 | 0.8430 |
| Symmetry imposed | C3 | C1 | C3 | C1 |
| Final particle images (no.) | 70,990 | 102,954 | 38,863 | 36,141 |
| Map resolution (Å) | 2.8 | 3.1 | 3.1 | 3.8 |
| FSC threshold | 0.143 | 0.143 | 0.143 | 0.143 |
| **Validation** |  |  |  |  |
| MolProbity score | 1.32 | 1.46 | 1.64 | 1.47 |
| Clash score | 1.85 | 2.54 | 2.28 | 0.9 |
| **Ramachandran plot** |  |  |  |  |
| Favored (%) | 94.52 | 94.1 | 95.27 | 95.65 |
| Allowed (%) | 99.61 | 5.32 | 99.43 | 4.35 |
| Outliers (%) | 0.39 | 0.57 | 0.57 | 0 |

**Table S2.**
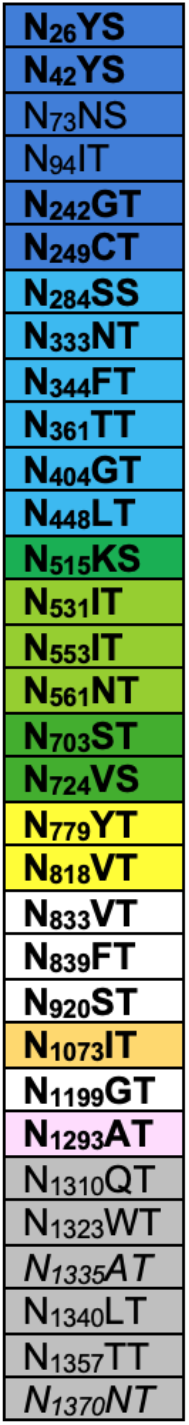
N-Linked glycosylation sequons in CCoV-HuPn-2018. Italic font indicates the absence of a glycosylation sequon. Glycans observed in either of the CCoV-HuPn 2P S cryo-EM maps are in bold. Glycans positions highlighted with the same color are located at the same CCoV-HuPn_2018 S domain or subunit. Color code: domain 0 (dark blue), domain A (light blue), domain B (light green), domain C (dark green), domain D (yellow) and S_2_ subunit (light orange, pin, white and grey).

**Table S3.** Biolayer interferometry kinetic parameters of B domains from HCoV-229E, CCoV-HuPN-2018 and TGEV immobilized at the surface of SA biosensors, to feline, canine and porcine APN orthologs and Fab 1AF10.

| Proteins | K<sub>D</sub> (nM) | k<sub>on</sub> (M<sup>-1</sup>s<sup>-1</sup>) | k<sub>off</sub> (s<sup>-1</sup>) |
| --- | --- | --- | --- |
| HCoV-229E Domain B human-APN | 9.3 | 6.6 x 10<sup>4</sup> | 6.1 x 10<sup>-4</sup> |
| CCoV-HuPn-2018 Domain B Feline-APN | 27 | 8 x 10<sup>4</sup> | 2.1 x 10<sup>-3</sup> |
| CCoV-HuPn-2018 Domain B Canine-APN | 1.0 | 1.7 x 10<sup>5</sup> | 1.8 x 10<sup>-4</sup> |
| CCoV-HuPn-2018 Domain B Porcine-APN | 41 | $3.5 \times 10^4$ | $1.4 \times 10^{-3}$ |
| CCoV-HuPn-2018 Domain B 1AF10 | 17 | $1.0 \times 10^5$ | $1.7 \times 10^{-3}$ |
| TGEV Domain B Feline-APN | 47 | $5.7 \times 10^4$ | $2.7 \times 10^{-3}$ |
| TGEV Domain B Canine-APN | 2.1 | $1.3 \times 10^5$ | $2.8 \times 10^{-4}$ |
| TGEV Domain B Pig-APN | 70 | $3.8 \times 10^4$ | $2.6 \times 10^{-3}$ |
| TGEV Domain B 1AF10 | 10 | $1.5 \times 10^5$ | $1.5 \times 10^{-3}$ |

## Methods

### Cell lines

Cell lines used in this study were obtained from ATCC: Human cells epithelial embryo (HEK293T, CRL-3216), *Felis catus* (CRFK, CCL-94), *Canis familiaris* epithelial kidney cells (MDCK, CCL-34), *Canis familiaris* tumor fibroblast cells (A-72, CRL-1542) and *Sus scrofa* pig testis fibroblast cells (ST, CRL-1746) or ThermoFisher Scientific: ExpiCHO cells and Expi293F™ cells. Cells were cultivated at 37°C, in an atmosphere of 5 % CO_2_ and with 130 rpm of agitation for suspension cells. None of the cell lines used were routinely tested for mycoplasma contamination.

### Plasmids

Genes used in this study were synthesized by GenScript, codon optimized for expression in mammalian cells, cloned into pcDNA3.1 (+) between KpnI and XhoI, in frame with a Kozak’s sequence to direct translation and with the signal peptide derived from the μ phosphatase: MGILPSPGMPALLSLVSLLSVLLMGCVAETGT (except for the full-length genes in which the original signal peptide was used). CCoV-HuPn-2018-S-2P corresponds to the sequence with the entry code: QVL91811.1 and includes residues 17 to 1387. To stabilize the spike in prefusion conformation, residues L (1140) and E (1141) were mutated to 2 P. Full-length wild-type S glycoproteins from HCoV-229E (AAK32191.1) residues: 1-1,155, CCoV-HuPn-2018 (QVL91811.1) residues 1-1,425 and TGEV (ABG89335.1) residues: 1-1,425, harbor C-terminal deletions of 18-, 23- and 23-residues, respectively, were used to pseudotype VSVΔG-luc. HCoV-229E, CCoV-HuPn-2018 and TGEV RBDs matching with the full-length sequences indicated above, include residues: 293 to 434, 539 to 671 and 522 to 665, respectively. Full-length APNs from human (NP_001141.2), feline (NP_001009252.2), canine (NP_001139506.1) and pig (AGX93258.1) orthologs comprise residues: 1-967, 1-967, 1-975 and 1-963, respectively. APNs wild-type ectodomains orthologs from human, feline, canine, pig, and human mutated (NWT) from the same sequence codes shown above, comprise residues: 66 to 967, 64 to 967, 71 to 975 and 62 to 963, respectively. The 1AF10 Fab light and heavy sequences were obtained from Reguera et al., 2012 and cloned separately into pcDNA3.1. Only the heavy chain was fused C-terminally with an 8 residues his-tag through a GGSGGS linker.

### Mutagenesis

Full-length wild-type human APN encoding plasmid was used as a template to create a glycosilation sequence at position N739 (NW<u>R</u> to NW<u>T</u> change) in a reaction mixture containing: 2X Kapa HiFi (Kapa biosystems) and the following primers, each at 10 μM: 5’-aataccaacaactgg<u>acc</u>gagatcc-3’ (underline is the codon change R to T) and 5’-atttctgaagtgaatgaagagg-3’ from Integrated DNA Technologies (IDT). The introduction of the desired mutation R to T was verified by sequencing purified plasmids by GENEWIZ. The plasmid harboring the mutation was further amplified and purified with EndoFree maxi kit (Qiagen) to be suitable for transfection into mammalian cells.

### Protein expression and purification

200 ml of ExpiCHO-S cells grown to a density of 6 x 10^6^/mL, were transfected with 640 μl of Expifectamine CHO reagent (ThermoFisher) and 200 μg of CCoV-HuPn-2018-S-2P following manufacturer’s recommendations. The day after transfection, feed and enhancer were added to the cells. Six days post-transfection, supernatants were clarified by centrifugation at 800 g for 10 minutes. Followed addition of 20 mM imidazole, 300 mM NaCl and 20 mM Tris-HCl pH 8.0, supernatants were further centrifuged at 14,000 g for 30 min and passed through a 1 mL His trap HP column (Cytiva) previously equilibrated with binding buffer (25 mM Tris pH 7.4 and 350 mM NaCl). CCoV-HuPn-2018-S-2P was eluted using a linear gradient of 500 mM imidazole.

To express B domains from CCoV-HuPn-2018, TGEV and HCoV-229E fused to an avidin and a histidine tag, 100 ml of Expi293F cells at 3 x 10^6^/mL were transiently transfected with 320 μl of Expifectamine and 100 μg of the respective plasmids, following the manufacturer’s indications. Four days post-transfection, supernatants were clarified by centrifugation at 800 g for 10 minutes, supplemented with 20 mM imidazole, 300 mM NaCl and 25 mM Tris-HCl pH 8.0, further centrifuged at 14,000 g for 30 min and passed through a 1 mL His trap HP column (Cytiva) previously equilibrated with binding buffer (25mM Tris pH 7.4 and 350mM NaCl). B domains were eluted using a linear gradient of 500 mM imidazole. Similar protocol was used to express and purify the Fab 1AF10 fused to 8 residues histidine tag except that 50 ml of Expi293F cells were transfected with a mixture containing: 50 μg of the individual plasmidsA encoding the light chain and heavy chain of the Fab and 160 μl of Expifectamine.

To express APN ectodomains from human, feline, canine and pig orthologs fused to Fc portion of human IgG, Expi293F cells were transiently transfected with the respective plasmids following the manufacturer’s protocols. Briefly, 50 ml of Expi293F cells at 3 x 10^6^/mL were transfected using 160 μl of Expifectamine and 50 μg of APN plasmid. Four days after transfection, supernatants were clarified by centrifugation at 800 g for 10 minutes, supplemented with 300 mM NaCl and 25 mM Tris-HCl pH 8.0, further centrifuged at 14,000 g for 30 min and passed through a 1 mL HiTrap Protein A HP column (Cytiva). Proteins were eluted using 0.1 M citric acid pH 3.0 in individual tubes containing 200 μl of 1 M Tris-HCl pH 9.0 to immediately neutralize the low pH needed for elution. Fractions containing the proteins were pooled and buffer exchanged to 25 mM Tris-HCl pH 8.0, 150 mM NaCl. To produce monomeric APNs, the Fc tag of all APN orthologs was removed using thrombin (Millipore Sigma) in a reaction mixture containing: 3 μg of thrombin/mg of APN-Fc, 20 mM Tris-HCl pH 8.0, 150 mM NaCl and 2.5 mM CaCl_2_ incubated overnight at room temperature. The reaction mixture was then loaded to a Protein A column to separate from uncleaved APN-Fc and the Fc tag. Monomeric APNs were further purified by size-exclusion chromatography (SEC) on a Superdex 200 column 10/300 GL (GE Life Sciences) previously equilibrated in 25 mM Tris pH 8.0 and 150 mM NaCl.

### Protein Biotinylation

B domains from HCoV-229E, TGEV and CCoV-HuPn-2018 were biotinylated using BirA biotin-protein ligase standard reaction kit (Avidity) following manufacturer’s protocol. In a typical reaction, 40μM of B domains were incubated overnight at 4°C with 2.5 μg of BirA enzyme in reaction mixtures containing 1X BiomixB, 1X BiomixA and 40 μM BIO200. Domains B were further purified by SEC using Superdex 75 increase 10/300 GL (GE LifeSciences) and concentrated using 10 kDa filters (Amicon).

### Biolayer interferometry

APNs binding measurements to B domains were performed using Biolayer interferometry. Biotinylated B domains from CCoV-HuPn-2018, TGEV and HCoV-229E were immobilized to strep-avidin (SA) biosensors, at 1 μg/mL in undiluted 10X kinetics buffer (Pall) to SA sensors that were pre-hydrated in water for at least 10 minutes and then equilibrated into 10X Kinetics Buffer (Pall). The RBDs were loaded to a level of 1 nm total shift. The loaded tips were then dipped into a dilution series of monomeric APN orthologs from human, feline, canine, and porcine or 1AF10 Fab in 10X Kinetics Buffer (Pall) starting at various concentrations for 300 seconds prior to 300 seconds dissociation in 10X Kinetics buffer for kinetics determination. The data were baseline subtracted and the plots fitted using the Pall FortéBio/Sartorius analysis software (v.12.0). Data were plotted in Graphpad Prism (v.9.0.2). These experiments were done side-by-side with two different batches of B domains and APN orthologs preparations. Competition BLI used 10X protein above measured K_D_ (for example, for CCOV-HuPn-2018 170nM of 1AF10 fab was used with a K_D_ of 17nM). CCoV-HuPn-2018 or TGEV biotinylated RBD was immobilized at 1 μg/mL in undiluted 10X kinetics buffer (Pall) to SA sensors that were prehydrated in water for at least 10 minutes and then equilibrated into 10X Kinetics Buffer (Pall). The RBDs were loaded to a level of 1 nm total shift. The loaded tips were then dipped first into 10X above K_D_ of 1AF10 fab diluted in 10X kinetics buffer, then into 1AF10 fab alone or 1AF10 fab plus 10X above K_D_ of with canine, feline, or porcine APN (Table 1). The data were baseline subtracted and fit using the Pall FortéBio/Sartorius analysis software (v.12.0). Data were plotted in Graphpad Prism (v.9.0.2).

### Pull-down assay

The interaction between APN orthologs and B domains was further analyzed performing a pull-down assay. Briefly, 200 μl (2mg) of magnetic beads Dynabeads™ Strep Isolation (ThermoFisher) were washed twice with 300 μL 1X TBS-T (2.4 g Tris-Base, 8 g NaCl, 0.1 % Tween20) using a magnetic stand (ThermoFisher) before coupling them with 200 μl of B domains in TBS at 0.125 ng/ml. The mixture beads/protein were well mixed and incubator at room temperature with constant gentle rotation. After 1 h incubation, beads were washed two times with 200 μL TBS-T before resuspending in a solution containing an excess of 10-20X of their calculated K_D_ using BLI, of the APN-Fc orthologs and allow binding for 1 hr at RT with gently rotation after which were washed two times with TBS-T. Flow through, last wash and beads were collected and analyzed in SDS-PAGE gel. Control assays consisted of uncoupled beads-APN-Fc.

### VSV-based pseudotyped virus production

CCoV-HuPn, HCoV-229E (AAK32191.1, P100E isolate) and TGEV S pseudotyped vesicular stomatitis virus (VSV), were generated as previously described (Tortorici et al., 2020, 2021). Briefly, HEK293T cells in DMEM supplemented with 19% FBS and 1% PenStrep and seeded in poly-D-lysine coated 10-cm dishes were transfected with a mixture of 24 μg of the corresponding plasmid encoding for: CCoV-HuPn S, TGEV S or HCoV-229 S, 60 μl Lipofectamine 2000 (Life Technologies) in 3 ml of Opti-MEM, following manufacturer’s instructions. After 5 h at 37°C, DMEM supplemented with 20% FBS and 2% PenStrep was added. Next day, cells were washed three times with DMEM and were transduced with VSVΔG-luc (Kaname et al., 2010). After 2 h, virus inoculum was removed and cells were washed five times with DMEM prior addition of DMEM supplemented with anti-VSV-G antibody [Il-mouse hybridoma supernatant diluted 1 to 25 (v/v), from CRL-2700, ATCC] to minimize parental background. After 18-24 h, supernatants containing pseudotyped VSVs were harvested, centrifuged at 2,000 x g for 5 minutes to remove cellular debris, filtered with a 0,45 μm membrane, concentrated 10 times using a 30 kDa cut off membrane (Amicon), aliquoted, and frozen at −80°C.

### Pseudotyped virus infections and neutralizations

For pseudotyped VSV infections and neutralizations, HEK293T cells were transfected with plasmids encoding for the different fulllength APN orthologs (flAPN) following the protocol described by (Eguia et al., 2021). Briefly, HEK293T cells at 90% confluency and seeded in poly-D-lysine coated 10-cm dishes were transfected with a mixture in Opti-MEM containing 8 μg of the corresponding plasmid encoding flAPN orthologs, 1 μg of the plasmid encoding full-length TMPRSS2 and 30 μl of Lipofectamine 2000 (Life Technologies) according to manufacturer’s instructions. After 5 h at 37°C, cells were trypsinized, seeded into poly-D-lysine coated clear bottom white walled 96-well plates at 40,000-50,000 cells/well and cultured overnight at 37°C. For infections, 20 μl of the corresponding pseudotyped VSV were mixed with 20 μl of DMEM and the mixture was added to the cells previously washed three times with DMEM. After 2 h at 37°C, 40 μl of DMEM was added and cells were further incubated ON at 37°C. For neutralizations, eleven 2-fold serial dilutions of Fab1AF10, APN ectodomains orthologs or sera, were prepared in DMEM. 20 μl of CCoV-HuPn-2018 S, HCoV-229E S or TGEV S pseudotyped VSV were added 1:1 (v/v) to each Fab1AF10, APN ectodomains or sera dilution and mixtures were incubated for 45-60 min at 37°C. After removing their media, transfected HEK293T cells were washed three times with DMEM and 40 μL of the mixture containing virus:Fab/APN ectodomains/sera were added. One hour later, 40 μL DMEM were added to the cells. After 17-20 h, 60 μL of One-Glo-EX substrate (Promega) were added to each well and incubated on a plate shaker in the dark. After 5-15 min incubation, plates were read on a Biotek plate reader. Measurements were done in duplicate with biological replicates. Relative luciferase units were plotted and normalized in Prism (GraphPad): cells alone without pseudovirus was defined as 0 % infection, and cells with virus only (no sera) was defined as 100 % infection. Most of the human sera was collected from prospective bone marrow donors in Seattle with approval from the Human Subjects Institutional Review Board in the 1980s and were stored in the Infectious Disease Sciences Biospecimen Repository at the Vaccine and Infectious Disease Division of the Fred Hutch Cancer Center. A few of the sera (the ones prefixed “FH”) are residual samples from Bloodworks Northwest that were collected from adults in Seattle.

### Western Blot

15 μl of pseudotyped VSV were mix with 4X SDS-PAGE loading buffer, run on a 4%–15% gradient Tris-Glycine Gel (BioRad) and transferred to PVDF membrane using the protocol mix molecular weight of the Trans-Blot Turbo System (BioRad). Membrane was blocked with 5% milk in TBS-T (20 mM Tris-HCl pH 8.0, 150 mM NaCL) supplemented with 0.05% Tween-20 at room temperature and with agitation. After 1 h, the fusion-peptide-specific S2S8 monoclonal antibody was added at 1:250 dilution and incubated ON at 4°C with agitation. Next day, the membrane was washed three times with TBS-T and an Alexa Fluor 680-conjugated goat antihuman secondary antibody (1:50,000 dilution, Jackson ImmunoResearch, 109-625-098) was added and incubated during 1 h at room temperature. Membrane was washed three times with TBS-T after which a LI-COR processor was used to develop the western blot.

### CryoEM sample preparation, data collection and data processing

Three microliters of recombinantly expressed and purified CCoV-HuPn-2018 at approximately 0.3 mg/ml were three times loaded onto freshly glow lacey grids covered with a thin layer of manually prior to plunge freezing using a vitrobot MarkIV (ThermoFisher Scientific) with a blot force of −1 and 2.5 sec blot time at 100% humidity and 21 °C. Data were acquired using the Leginon software (Suloway et al., 2005) to control a FEI Titan Krios transmission electron microscope equipped with a Gatan K3 direct detector and operated at 300 kV with a Gatan Quantum GIF energy filter. The dose rate was adjusted to 3.75 counts/super-resolution pixel/s, and each movie was acquired in 75 frames of 40 ms with a pixel size of 0.843 Å and a defocus range comprised between −0.8 and −2.0 μm. Movie frame alignment, estimation of the microscope contrast-transfer function parameters, particle picking and extraction (with a sampled pixel size of 1.686 Å and box size of 320 pixels^2^) were carried out using Warp (Tegunov and Cramer, 2019). Two rounds of reference-free 2D classification were performed using cryoSPARC (Punjani et al., 2017) to select well-defined particle images. Subsequently, one round of 3D classification with 50 iterations, using ab initio as a reference model (angular sampling 7.5 ° for 25 iterations and 1.8 ° with local search for 25 iterations) was carried out using Relion (Zivanov et al., 2018) without imposing symmetry. 3D refinements were carried out using non-uniform refinement in cryoSPARC (Punjani et al., 2020). Particle images were subjected to Bayesian polishing (Zivanov et al., 2019) using Relion during which particles were re-extracted with a box size of 512 pixels at a pixel size of 0.843 Å which was followed by another round of non-uniform refinement in cryoSPARC followed by per-particle defocus refinement and again non-uniform refinement. To improve the density of the domain 0, particles were symmetry-expanded and subjected to a Relion focus 3D classification without refining angles and shifts using a soft mask encompassing domain 0. Local resolution estimation, filtering, and sharpening were carried out using CryoSPARC. Reported resolutions are based on the gold-standard Fourier shell correlation (FSC) of 0.143 criterion (Rosenthal and Henderson, 2003) and Fourier shell correlation curves were corrected for the effects of soft masking by high-resolution noise substitution (Chen et al., 2013).

### CryoEM model building and analysis

UCSF Chimera (Pettersen et al., 2004) and Coot (Emsley et al., 2010) were used to fit atomic models (PDB 5SZS) into the cryoEM maps and Domain 0 was manually built. Models were refined and rebuilt into the maps using Coot and Rosetta (Frenz et al., 2019; Wang et al., 2016). Model validation was done using (Chen et al., 2010) and Privateer (Agirre et al., 2015). Figures were generated using UCSF ChimeraX (Goddard et al., 2018). Topology diagrams were generated using the program TopDraw (Bond, 2003).

### Assessment of human *ANPEP* diversity

We assessed human diversity at or near the NWT motif (NWR in human; R741) in Gnomad (https://gnomad.broadinstitute.org/gene/ENSG00000166825). Observations in approximately 125,000 humans included one heterozygous N738K, one heterozygous R741G and one heterozygous I743N. No rare defects in *ANPEP* are described in OMIM (https://www.omim.org/entry/151530)

## Notes

### Competing Interest Statement

The authors have declared no competing interest.

